# The HCP 7T Retinotopy Dataset: Description and pRF Analysis

**DOI:** 10.1101/308247

**Authors:** Noah C. Benson, Keith W. Jamison, Michael J. Arcaro, An Vu, Matthew F. Glasser, Timothy S. Coalson, David C. Van Essen, Essa Yacoub, Kamil Ugurbil, Jonathan Winawer, Kendrick Kay

## Abstract

About a quarter of human cerebral cortex is dedicated mainly to visual processing. The large-scale organization of visual cortex can be measured with functional magnetic resonance imaging (fMRI) while subjects view spatially modulated visual stimuli, also known as ‘retinotopic mapping’. One of the datasets collected by the Human Connectome Project (HCP) involved ultra-high-field (7 Tesla) fMRI retinotopic mapping in 181 healthy young adults (1.6-mm resolution), yielding the largest freely available collection of retinotopy data. Here, we describe the experimental paradigm and the results of model-based analysis of the fMRI data. These results provide estimates of population receptive field position and size. Our analyses include both results from individual subjects as well as results obtained by averaging fMRI time-series across subjects at each cortical and subcortical location and then fitting models. Both the group-average and individual-subject results reveal robust signals across much of the brain, including occipital, temporal, parietal, and frontal cortex as well as subcortical areas. The group-average results agree well with previously published parcellations of visual areas. In addition, split-half analyses show strong within-subject reliability, further demonstrating the high quality of the data. We make publicly available the analysis results for individual subjects and the group average, as well as associated stimuli and analysis code. These resources provide an opportunity for studying fine-scale individual variability in cortical and subcortical organization and the properties of high-resolution fMRI. In addition, they provide a set of observations that can be compared with other HCP measures acquired in these same participants.

## Introduction

The central nervous system maps sensory inputs onto topographically organized representations. In the field of vision, researchers have successfully exploited functional magnetic resonance imaging (fMRI) to noninvasively measure visual field representations (‘retinotopy’) in the living human brain (DeYoe et al., 1996; Engel, Glover, & Wandell, 1997; Engel et al., 1994; Sereno et al., 1995). These efforts enable parcellation of visual cortex into distinct maps of the visual field, thereby laying the foundation for detailed investigations of the properties of visual cortex (*parcellation references*: Abdollahi et al., 2014; Benson, Butt, Brainard, & Aguirre, 2014; Wang, Mruczek, Arcaro, & Kastner, 2015; *review references*: Silver & Kastner, 2009; Tootell, Dale, Sereno, & Malach, 1996; Wandell & Winawer, 2011; Wandell, Dumoulin, & Brewer, 2007).

One of the datasets acquired by the Human Connectome Project (HCP) (Ugurbil et al., 2013; Van Essen et al., 2013) was a 7T fMRI retinotopy experiment. This experiment, conducted in 181 healthy young adults, involved carefully designed stimuli and a substantial amount of fMRI data (30 minutes, 1,800 time points) acquired at high spatial and temporal resolution (1.6-mm isotropic voxels, 1-second sampling). Although retinotopy is routinely measured in small groups of subjects by individual laboratories in support of various research projects, to date there has not been a large publicly available set of retinotopic measurements.

In this paper, we describe the design of the retinotopy experiment and demonstrate the analyses that we have performed on the fMRI data. We adopt a model-based analysis approach in which a computationally intensive nonlinear optimization is performed to determine parameters of a population receptive field (pRF) model (Dumoulin & Wandell, 2008; Kay, Winawer, Mezer, & Wandell, 2013; Wandell & Winawer, 2015). The results include estimates of pRF position (angle and eccentricity) and pRF size for each ‘grayordinate’ (cortical surface vertex or subcortical voxel), and can be used to define retinotopic maps in the brain. We show that the HCP retinotopy data provide high-quality pRF results in many parts of occipital, temporal, parietal, and frontal cortex. We make freely available these pRF results, as well as associated stimuli and analysis code, at an Open Science Framework web site (https://osf.io/bw9ec/). The pRF results are also accessible via the BALSA database (https://balsa.wustl.edu/study/show/9Zkk; Van Essen et al., 2017), downloadable as ‘scene files’ that can be visualized using Connectome Workbench software (see Appendix). The neuroscience community at large can now exploit these resources for a variety of purposes, such as developing normative models, mapping new brain areas, analyzing connectomics, characterizing individual differences, and comparing with other suitably aligned datasets (either published or ongoing).

## Results

Here we present a summary of the data quality and example results from the HCP 7T Retinotopy Dataset. The stimuli and analyses are detailed in the Methods and are described here very briefly. Each of 181 subjects participated in six 5-minute pRF mapping runs. The stimuli comprised colorful object textures windowed through slowly moving apertures (Figure 1A). The colorful object textures were used because they produce high signal-to-noise ratio in higher-level visual areas. The apertures were clockwise or counterclockwise rotating wedges, expanding or contracting rings, or bars that swept across the visual field in several directions (Figure 1B).

**Figure 1.**
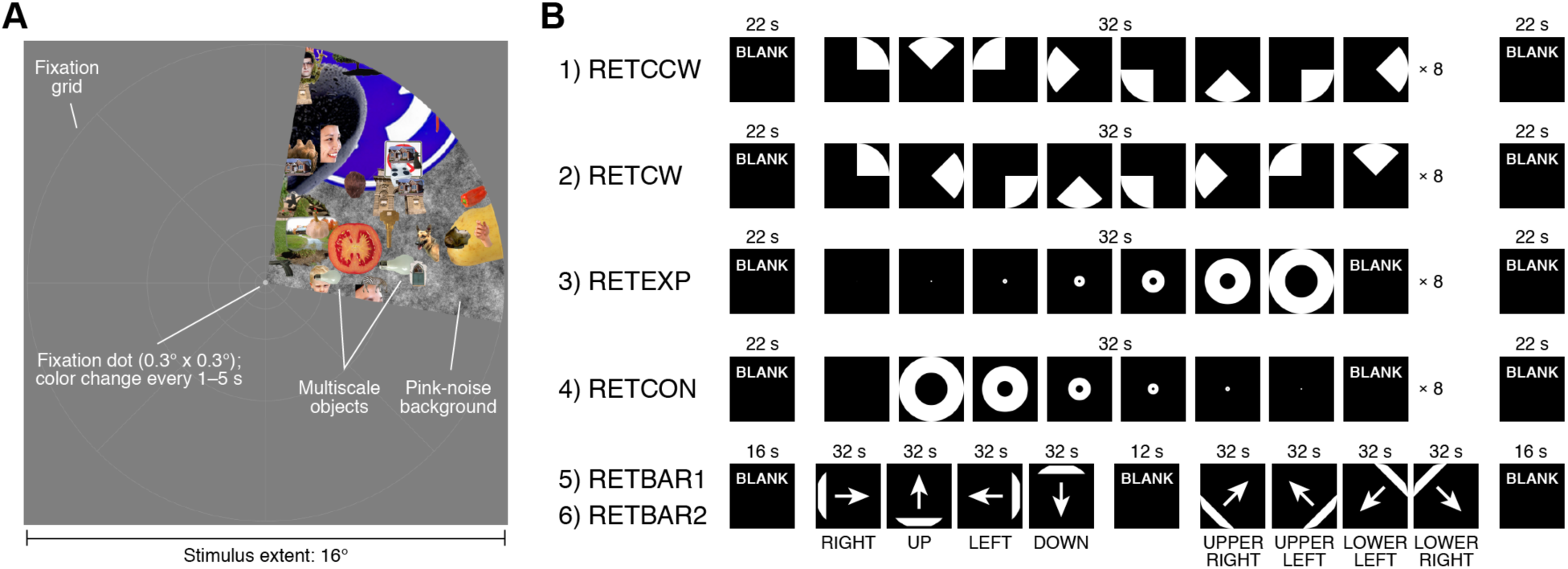
Schematic of experiment. (A) Example stimulus frame. The stimulus consisted of a dynamic colorful texture (composed of objects at multiple scales placed on a pink-noise background) presented within a slowly moving aperture. The aperture and texture were updated at 15 Hz. Subjects were instructed to fixate on a small fixation dot and to press a button whenever its color changed. A fixation grid was provided to aid fixation. (B) Run design. Six 300-s runs were acquired. The temporal structure of the runs is depicted. The first two runs involved a rotating wedge (RETCCW, RETCW), the second two runs involved an expanding or contracting ring (RETEXP, RETCON), and the last two runs involved a moving bar (RETBAR1, RETBAR2).

The resource we provide with this paper is a large set of population receptive field (pRF) model solutions. We define the pRF as the region of the visual field within which a visual stimulus elicits an increase in response from the pooled neural activity reflected in fMRI measurements, and can be summarized by the pRF’s angle, eccentricity, and size (Figure 2A). The total dataset consists of 181 individual subjects and 3 group averages. The 3 group averages reflect two split-halves of the subjects as well as all 181 subjects. For each of the 181 individuals and the 3 group averages, we solved 3 sets of models: one from the concatenation of all 6 runs (300 seconds per run, 1,800 time points), one from the first half of each run (150 seconds per run, 900 time points), and one from the second half of each run (150 seconds per run, 900 time points). For each subject or group average and for each of the 3 types of model fits, we obtained model solutions for the 91,282 cortical vertices and subcortical voxels (‘grayordinates’ spaced on average 2 mm apart). Each model solution yielded 6 numbers: angle, eccentricity, pRF size, and gain describing the pRF model, variance explained by the model, and mean BOLD signal. Therefore in total, the pRF model solutions that we provide consist of 184 subjects (181 individuals and 3 group averages) × 91,282 grayordinates × 3 model fits × 6 quantities (Figure 2B). Individual subjects are referred to as S1–S181, the two split-half group averages are referred to as S182 and S183, and the full group average is referred to as S184.

**Figure 2.**
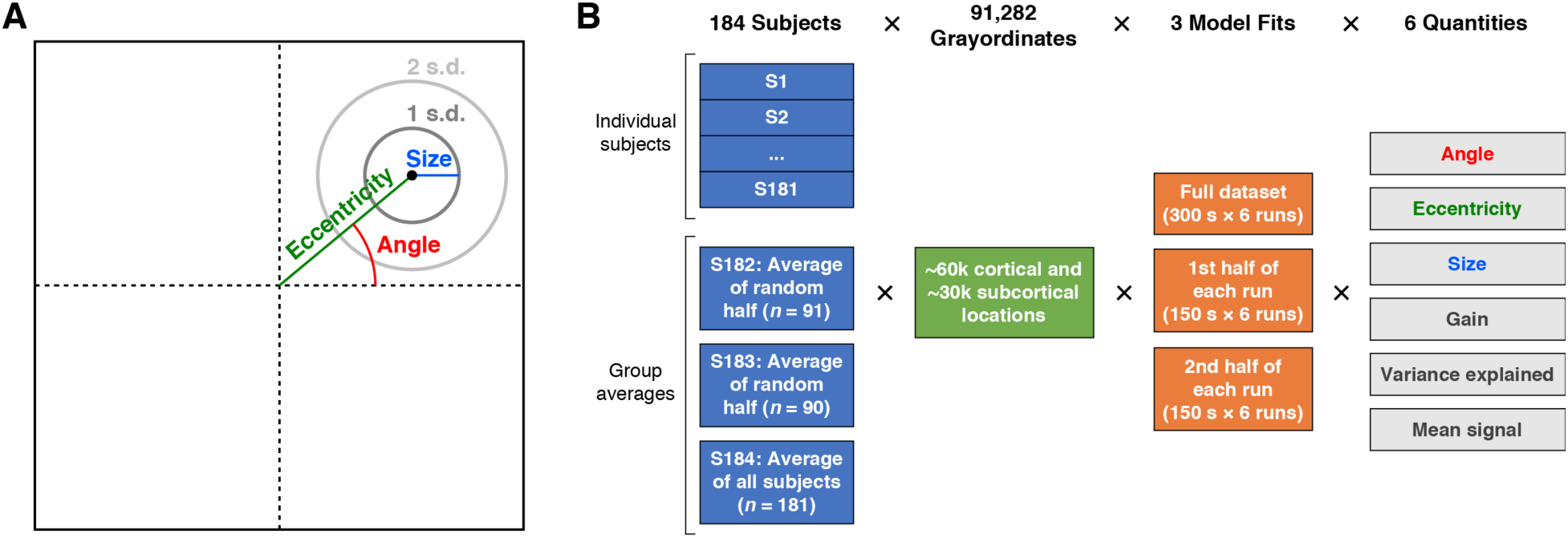
pRF model solutions provided in this resource. (A) pRF parameters. Each pRF is described by a 2D Gaussian. Angle is the rotation of the center of the Gaussian with respect to the positive *x*-axis. Eccentricity is the distance between the center of gaze and the center of the Gaussian. Size is defined as one standard deviation (s.d.) of the Gaussian. Angle is in units of degrees of polar angle, whereas eccentricity and size are in units of degrees of visual angle. (B) pRF model solutions. We solved pRF models for 181 individual subjects and 3 group-average pseudo-subjects (the average of split-halves of the subjects or of all subjects). For each individual subject (S1–S181) and each group average (S182–S184), 3 types of models were fit: one reflecting the complete set of runs and two reflecting split-halves of the runs. Model fits were obtained independently for each of 91,282 grayordinates, yielding 6 quantities. The total dimensions of the pRF model solutions are 184 subjects × 91,282 grayordinates × 3 model fits × 6 quantities.

The particular form of the pRF model we employed assumes that each grayordinate’s pRF is a 2D isotropic Gaussian and that contrast within the pRF is summed sublinearly according to a static power-law nonlinearity with exponent 0.05 (Kay et al., 2013a). This can be expressed formally as

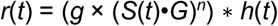

where *r*(*t*) is the predicted stimulus-related time series, *g* is a gain parameter, *S*(*t*) is the stimulus aperture at time *t, G* is the 2D isotropic Gaussian, *n* is an exponent parameter (*n* = 0.05), and *h*(*t*) is a canonical HRF. The sub-additive exponent was used to obtain more accurate pRF solutions, but since it was fixed for all models we do not analyze it further. Note that the pRF sizes that we report take into account the effects of the sub-additive exponent (see Methods for details).

### Distinctions between visual responsivity, spatial selectivity, retinotopic organization, and retinotopic maps

The pRF model solutions can be used to make different types of inferences regarding visual response properties. First, if the pRF model successfully explains variance in the time-series data for a grayordinate, this indicates that the grayordinate is *visually responsive*, but does not by itself imply spatial selectivity. For example, if a grayordinate responds with equal strength to a stimulus presented anywhere in the visual field, variance in its time-series data can be explained by a pRF model that has very large spatial extent. Second, to be considered *spatially selective*, a grayordinate must not only be visually responsive but also exhibit larger responses to stimuli in some locations compared to others. To help assess spatial selectivity in the HCP dataset, we fit each grayordinate time series with a simple ON/OFF model that is sensitive only to the presence or absence of the stimulus, and compare the variance explained by this model to that explained by the full pRF model (see Supplementary Figure 1). Visual responsivity and spatial selectivity are properties of a single grayordinate. A third type of inference is *retinotopic organization*. This is a stronger claim that describes spatial selectivity at a larger scale: retinotopic organization implies not only that single brain locations are spatially selective, but also that adjacent brain locations respond to nearby locations in the visual field, thereby producing smooth progressions of polar angle and/or eccentricity. In principle, a brain region might be spatially selective but not retinotopic if the spatial tuning of nearby brain locations is haphazard. In practice, this seems to be uncommon. Finally, a *retinotopic map*, or *visual area*, is generally considered to be a region of the brain that contains a representation of all or most of the contralateral visual hemifield in each hemisphere. In this paper, we make observations regarding retinotopic organization in the HCP dataset but do not attempt to resolve various ongoing controversies regarding human retinotopic maps (see Discussion).

### Group-average results

#### Cortical data

We first summarize pRF model solutions from group average S184 (which reflects all 181 individual subjects). Group-average results were obtained by taking the time-series data from individual subjects (aligned using MSMAll to HCP’s average cortical surface space fs_LR; see Methods), computing the across-subject average of the time-series data observed at each grayordinate, and then fitting a pRF model to the time-series data at each grayordinate. For visualization, we map the results from fs_LR space to *fsaverage* space and plot the results on the *fsaverage* surface that has been inflated, spherized, and orthographically projected to a plane (Figure 3). We also provide visualizations of the results on the inflated *fsaverage* surface using dynamic rotating movies (Supplementary Movies 1–12).

**Figure 3.**
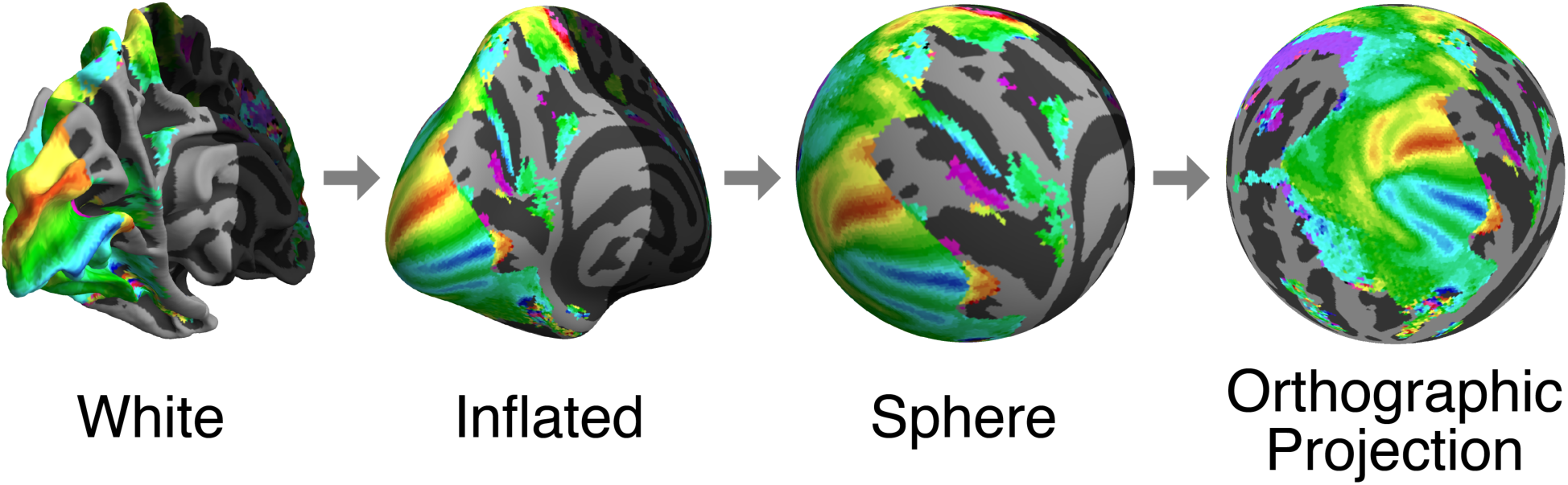
Cortical surface visualization. Cortical surfaces are inflated, warped to a sphere, and rotated with shading discarded to render as an orthographic projection. The regions of the first two surfaces (white, inflated) that are not visible in the final view are darkened. For this schematic, we depict the thresholded group-average angle results (see Figure 4) to provide a visual reference across the transformations.

The effect of averaging the time-series data across subjects differs across the cortex, depending on how well pRF parameters match between subjects given the MSMAll alignment. Prior work has shown that the V1–V3 maps have highly regular topography and are well aligned to measures of anatomy, such as surface curvature (Benson et al., 2014; 2012; Hinds et al., 2008) and myelination (Abdollahi et al., 2014), and to measures of function such as resting-state connectivity (Bock et al., 2015; Raemaekers et al., 2014). Therefore, these maps are likely to be well aligned across subjects, and averaging will preserve many of the features found in the maps of individual subjects. In particular, the angle and eccentricity maps show clear and expected patterns in V1–V3 (Figure 4, second and third columns), and the variance explained is greater than 75% (cyan regions in the fifth column of Figure 4). As expected, from the lower to upper bank of the calcarine sulcus, there is a smooth progression from the upper vertical meridian through the contralateral horizontal meridian to the lower vertical meridian (blue-cyan-green-yellow-red sweep in the angle colormaps). The angle map reverses at the lips of the calcarine sulcus, with mirror-reversed and approximately quarter-field representations in the bordering dorsal and ventral V2 maps and dorsal and ventral V3 maps. As expected, the eccentricity map is in register across V1–V3, progressing from foveal to peripheral representations from near the occipital pole towards medial and anterior directions (blue-magenta-red-yellow-green progression in the eccentricity colormap). The pRF size map has some of the same features of the eccentricity map, exhibiting smaller sizes near the occipital pole and larger sizes in the mid-peripheral regions of V1–V3. However, in the more peripheral portions of the maps, the size estimates are smaller than predicted from eccentricity due to stimulus edge effects (blue rim around the anterior/medial edge of the V1–V3 maps; see Discussion).

**Figure 4.**
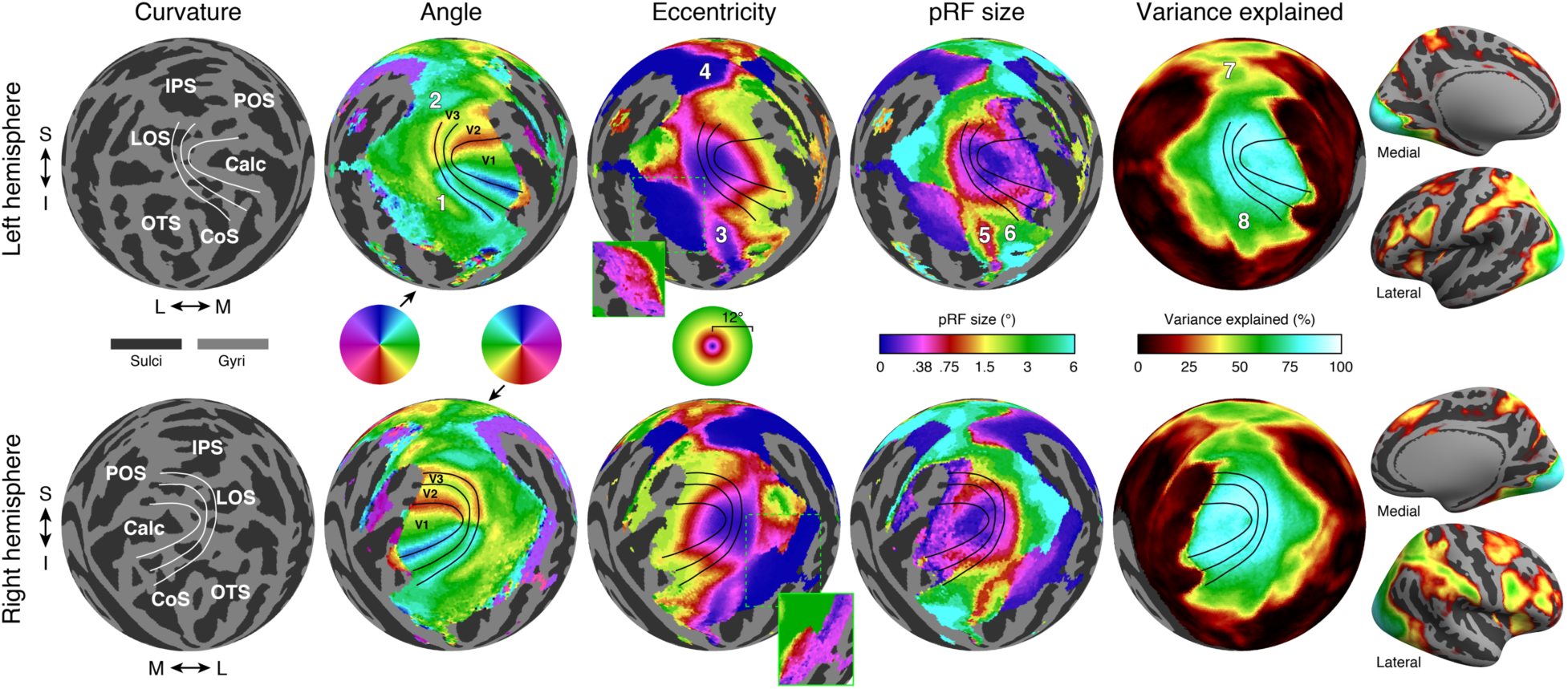
Group-average results. The pRF model solutions are mapped from fs_LR space to *fsaverage* using nearest-neighbor interpolation and then visualized (see Methods). Here we visualize results for group average S184 (which reflects all 181 individual subjects) in occipital cortex using an orthographic projection (see Figure 3). The first column shows the thresholded *fsaverage* curvature. White lines are hand-drawn borders of V1, V2, and V3 based on the angle results. Labels indicate several major posterior sulci. The second through fourth columns show angle, eccentricity, and pRF size maps (with areal boundaries now shown in black). These maps are thresholded at 9.8% variance explained (see Methods). In the eccentricity maps, the insets marked with green show the same results but with the entire color range corresponding to 0–0.5°— this demonstrates that the large uniform swath of blue in the main figure actually has gradients of near-foveal eccentricities. The fifth column shows variance explained. Finally, the images on the right show thresholded variance explained on inflated left and right surfaces, demonstrating the existence of robust signals in other parts of cortex. Labels: S = superior, I = inferior, M = medial, L = lateral, IPS = intraparietal sulcus, LOS = lateral occipital sulcus, POS = parieto-occipital sulcus, Calc = calcarine sulcus, OTS = occipitotemporal sulcus, CoS = collateral sulcus. In the top panels, the numbers 1–8 indicate features of the parameter maps and are discussed in the text. Distinct numbers are used in each parameter map although some locations are the same or nearly the same (locations 4 and 7; locations 3, 5, and 8).

In cortical locations where pRF parameters are variable across subjects (even after registration using MSMAll), the group-average results will preserve less of the detail from individual subjects. Nonetheless, there is a large amount of structure in the group-average results beyond V1–V3, and some clear patterns are evident. The angle maps show the expected progression from upper to lower field ventral to V3, and from lower to upper field dorsal to V3 (locations 1 and 2 in Figure 4, top row), consistent with measurements of ventral (Kastner et al., 2001; McKeefry, Watson, Frackowiak, Fong, & Zeki, 1997; Wade, Brewer, Rieger, & Wandell, 2002) and dorsal (Press, Brewer, Dougherty, Wade, & Wandell, 2001; Tootell et al., 1997) occipital cortex. The eccentricity map also shows clear large-scale organization throughout large expanses of parietal and temporal cortex. One feature of the eccentricity maps is multiple distinct, foveal representations: in addition to the foveal representation in V1–V3 at the occipital pole, the eccentricity maps show distinct foveal representations in ventral temporal cortex and parietal cortex (locations 3 and 4 in Figure 4), consistent with many prior studies (Swisher, Halko, Merabet, McMains, & Somers, 2007; Tootell et al., 1997; Wade et al., 2002; Wandell, Brewer, & Dougherty, 2005). Near both of these distinct foveal representations, there are foveal to peripheral progressions along the lateral to medial direction.

The pRF size map also shows a variety of large-scale patterns. In ventral temporal cortex, there is a small-to-large size gradient from the fusiform gyrus to the collateral sulcus (locations 5 and 6 in Figure 4). These regions roughly correspond to the locations of face-selective and place-selective cortex (Epstein & Kanwisher, 1998; Grill-Spector & Weiner, 2014; Kanwisher, McDermott, & Chun, 1997). More generally, pRF sizes tend to be larger outside V1–V3, as expected from both single-unit and fMRI measurements (Dumoulin & Wandell, 2008; Maunsell & Van Essen, 1983; Smith, Singh, Williams, & Greenlee, 2001; Tootell et al., 1997). Finally, the variance explained map shows that robust signals occur not only within V1–V3 but also in higher-level areas. Variance explained is above 50% in several regions ventral, lateral, and dorsal to the V1–V3 maps, including much of ventral temporal cortex and the intraparietal sulcus (locations 7 and 8 in Figure 4). Furthermore, for nearly all the cortical locations that survive the variance explained threshold, pRF model parameters are highly reliable. This is confirmed by two types of split-half analysis: First, the data averaged across all 181 subjects (S184) were split into two halves by time, with one dataset comprising time series from the first half of each of the six runs, and a second dataset comprising time series from the second half of each of the six runs. Second, the data were split into two halves by subject, with one dataset reflecting averaging time-series data across 91 subjects (S182), and a second dataset reflecting averaging time-series data across the remaining 90 subjects (S183). Both split-half analyses indicate high reliability of pRF parameters (results not shown; pRF model solutions available online).

#### Relationship to cortical parcellations

Many features of the group-average results are in good agreement with recently published parcellations of visual areas, particularly near the posterior occipital pole (Figure 5). We compare the group-average pRF results to two atlases made using different methods: the Wang et al. (Wang et al., 2015) maximum probability atlas and the Glasser et al. (Glasser, Coalson, et al., 2016a) multimodal parcellation of cortex. The Wang et al. maximum probability atlas includes 25 regions of interest (ROIs) per hemisphere whose boundaries are derived from the anatomically-aligned overlap of manually labeled visual areas in individual subjects. Ten of these in posterior cortex are clearly aligned with expected features of the angle maps: V1v/V1d, V2v/V2d, V3v/V3d, V3A, V3B, LO-1, and hV4. In each of these 10 ROIs, one or more borders lie on an angle reversal. For example, the V1d/V2d border lies on a lower-field angle reversal, and the V1v/V2v border lies on an upper-field angle reversal. The agreement between the retinotopic features of the HCP dataset and the borders from the Wang et al. atlas is remarkably good despite the atlas reflecting different subjects and different experimental and analysis methods. Other maps such as LO-2, TO-1/2, IPS maps, VO-1/2, and PHC-1/2 show contralateral representations but not clear progressions of angle, in part due to blurring from group averaging (i.e., averaging cortical locations from different subjects that represent different portions of the visual field). The Glasser et al. parcellation was generated using a semi-automated, supervised approach applied to multimodal neuroimaging data (representing architecture, connectivity, function, and visuotopic organization). For the Glasser et al. parcellation, several areas are well aligned with features of the retinotopic maps, particularly V1, V2, V3, V4, and V3A. In several map clusters in the Wang et al. atlas, there are clear eccentricity gradients: the IPS0–2 maps show a clear foveal-to-peripheral gradient along the medial-to-lateral direction, as do the V1–V3 maps and the VO-1/2 maps. In the Glasser et al. atlas, several areas fall within iso-eccentricity regions. For example, the areas PH and TE2p are clearly foveal, whereas the adjacent fusiform face complex (FFC) is more peripheral, with a sharp change in eccentricity along the border between these areas. Consistent with our definitions in the previous sections, the eccentricity gradients indicate clear retinotopic organization in these anterior areas; however, we do not try to draw conclusions in this paper regarding whether there are complete maps of the visual field or the most appropriate way to parcellate cortex into distinct areas.

**Figure 5.**
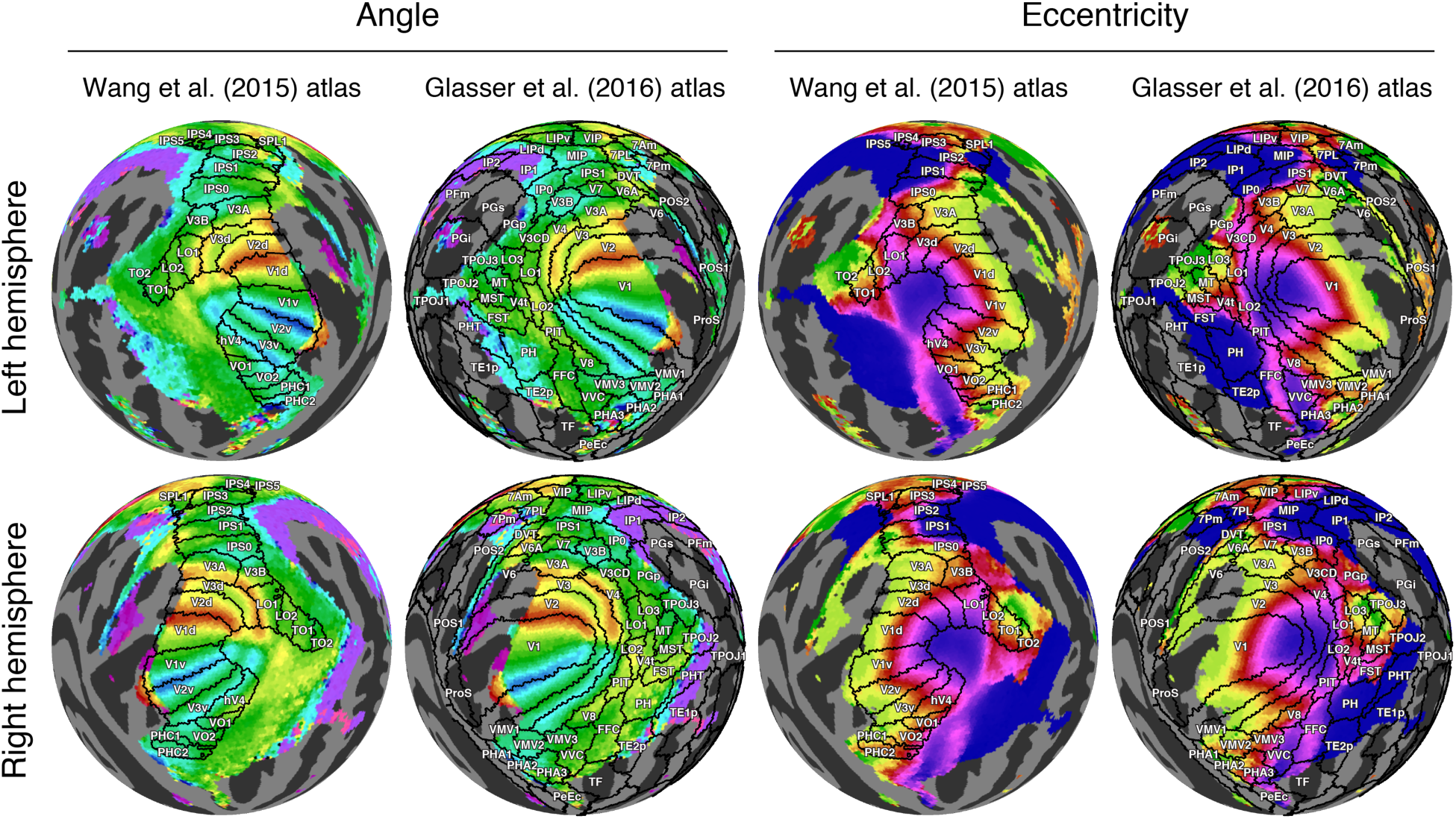
Relationship between group-average results and cortical parcellations. The angle and eccentricity maps from the group-average dataset (S184) are re-plotted from Figure 4 (same color scales). Superimposed on the maps are regions of interest from the maximum probability atlas of Wang et al. (Wang et al., 2015) and cortical parcellations from Glasser et al. (Glasser, Coalson, et al., 2016a).

#### Subcortical data

The HCP 7T Retinotopy Dataset includes subcortical results in addition to cortical results. Several subcortical nuclei have retinotopic maps that have been previously measured using fMRI (Arcaro, Pinsk, & Kastner, 2015; Cotton & Smith, 2007; DeSimone, Viviano, & Schneider, 2015; Katyal, Zughni, Greene, & Ress, 2010; Schneider & Kastner, 2005; Schneider, Richter, & Kastner, 2004). The subcortical fMRI data were aligned using FNIRT nonlinear volume registration based on T1-weighted image intensities (Glasser et al., 2013). In contrast to cortex, subcortical structures are not easily represented as 2D surfaces, and hence it is more difficult to visualize complete maps. Nonetheless, slices through subcortical structures reveal clear, high-quality pRF model solutions in the group-average dataset (Figure 6). In particular, we see expected structure in visual nuclei such as the lateral geniculate nucleus (LGN), superior colliculus (SC), and ventral pulvinar (vPul1/2). Within these regions, there are clear representations of the contralateral visual field. As expected, the visual field maps of the LGN and pulvinar are both inverted with smooth progressions from the upper visual field located ventrally to the lower visual field located dorsally. In the superior colliculus, there is a smooth progression from the upper visual field (anterior and medial) to the lower visual field (posterior and lateral).

**Figure 6.**
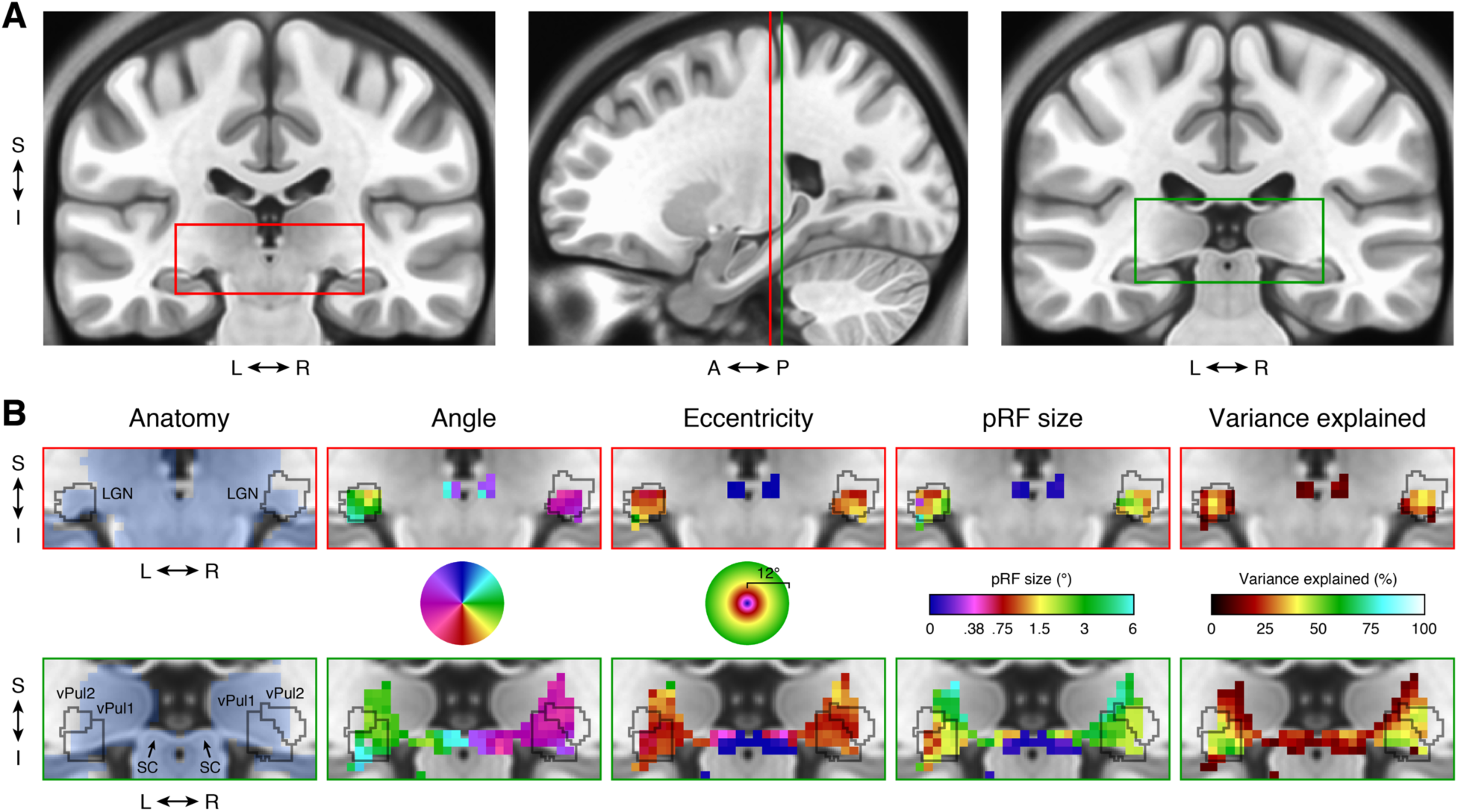
Subcortical results. (A) Anatomical location. The two coronal slices (*y* = 25, far left; *y* = 30, far right) show the MNI average anatomy (ICBM 152 nonlinear symmetric atlas 2009b, 0.5-mm resolution). The red and green rectangles mark the regions detailed in panel B. Vertical lines on the sagittal slice (*x* = 23) indicate the locations of the two coronal slices. (B) pRF results. The upper row highlights the left and right LGN and the lower row highlights the pulvinar and superior colliculus. Outlines of the LGN and ventral pulvinar (vPul1/2) are taken from Arcaro et al., 2015. All pRF results are from the group-average dataset (S184) and are thresholded at 9.8% variance explained, as in Figure 4. Colormaps are identical to those in Figure 4 except that only the left-hemisphere angle colormap is used. The blue shading in the anatomy column indicates voxels that are included in the CIFTI subcortical data mask.

### Individual-subject results

In addition to group-average results, we also computed pRF model solutions for the 181 individual subjects. We summarize results in several ways, including quantifying the amount of variance explained by the pRF model, inspecting maps in individual subjects, and assessing within-subject reliability of pRF parameters. These analyses reveal that overall data quality is high.

#### Variance explained

We quantified variance explained by the pRF model within atlas-defined ROIs. We defined one ROI as the union of the 50 maps found in the Wang et al. maximum probability atlas (25 maps per hemisphere) and a second ROI as the union of the V1–V3 maps from the same atlas (Wang et al., 2015). The V1–V3 ROI is a subset of the larger ROI. Because these ROIs are defined based on group-average anatomy, they do not necessarily conform to each individual subject’s retinotopic maps, but they provide a simple objective method for region definition. Within the union of the 50 maps, we computed for each subject the median variance explained across grayordinates, yielding one number per subject. The median of this number across the 181 subjects was 17% (Figure 7A). Within just the V1–V3 maps, the median of the median variance explained was substantially higher, at 44%. For comparison, we estimate that for grayordinates not sensitive to the experimental paradigm, the variance explained by the pRF model is less than 1%. This can be seen by inspecting the large peak in the histogram of variance explained across all grayordinates from all individual subjects (Figure 7C).

**Figure 7.**
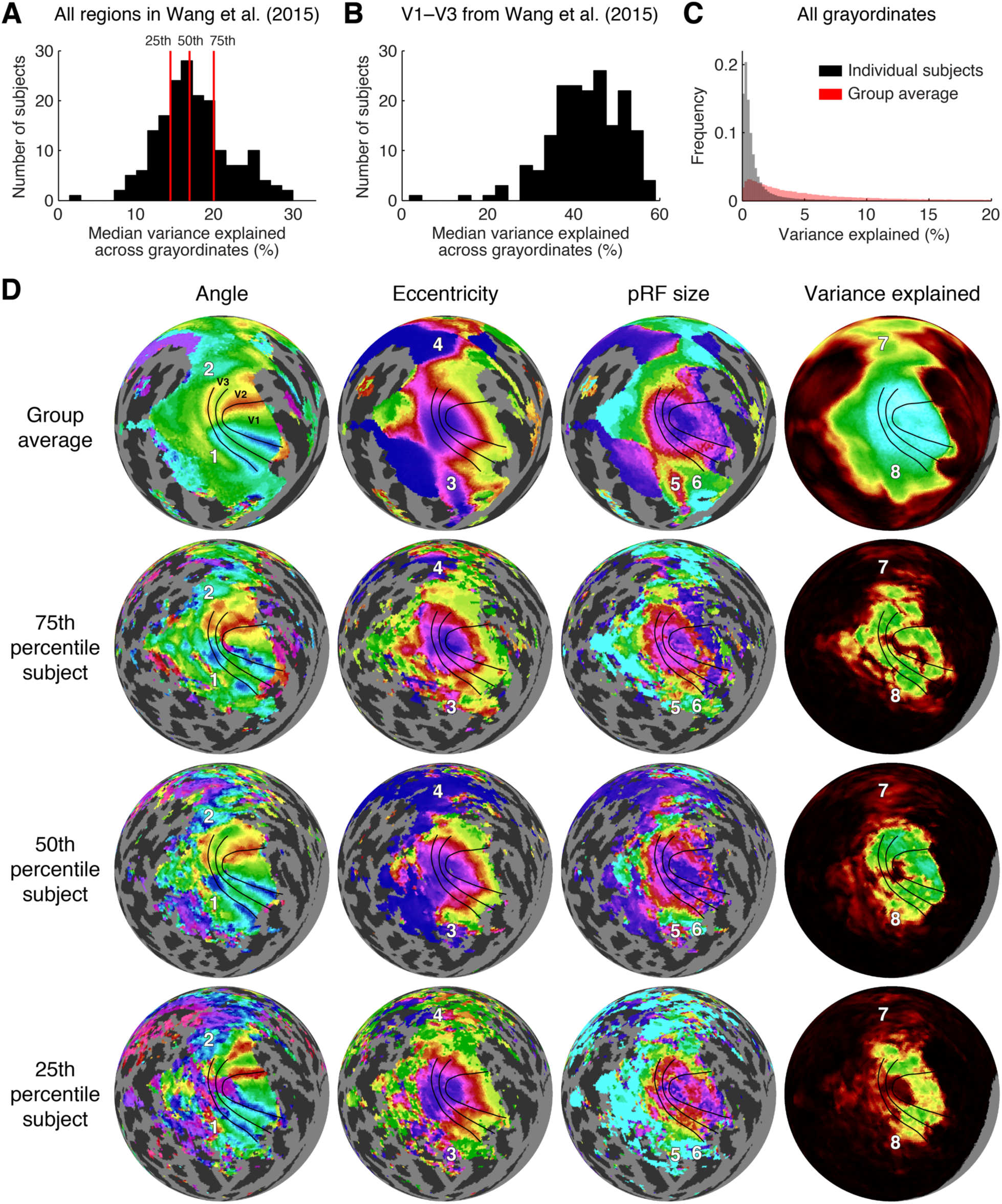
Individual-subject results. (A) Variance explained within all regions of the Wang et al. maximum probability atlas (Wang et al., 2015). For each subject, we computed the median variance explained across grayordinates located within the union of all regions in both hemispheres of the Wang et al. atlas. The histogram shows the distribution of this value across the 181 subjects. The subjects at the 25^th^, 50^th^, and 75^th^ percentiles are indicated by red lines. (B) Variance explained within V1–V3 of the Wang et al. atlas. (C) Histogram of variance explained across all grayordinates in individual subjects S1–S181 and in the group average S184 (bin size 0.2%; histogram counts normalized to sum to 1 in each plot). (D) Maps of pRF parameters (left hemisphere). We re-plot the group-average results from Figure 4 (S184) and show individual-subject results for the three subjects indicated in panel A (S43, S11, and S124, corresponding to the 25^th^, 50^th^, and 75^th^ percentiles, respectively; corresponding HCP IDs 164131, 115017, and 536647). Angle, eccentricity, pRF size, and variance explained results are plotted as in Figure 4 (with the same color scales), except that the variance explained threshold used for individual subjects is 2.2% (see Methods). For reference, we show on each map the same V1–V3 boundary lines determined from group-average results in Figure 4 as well as the same numbered locations (1–8) that mark features of the parameter maps.

#### Cortical maps

For map visualization, we selected three representative subjects: the subjects at the 25^th^, 50^th^, and 75^th^ percentiles with respect to median variance explained across regions in the Wang et al. atlas (see red lines in Figure 7A). For simplicity we show only the left hemisphere, and we re-plot the group-average results for comparison. The three depicted subjects have clear retinotopic maps in occipital cortex, as seen in the angle and eccentricity results (Figure 7C). In each subject, the angle maps reveal the boundaries of V1–V3, and the eccentricity maps are in register across visual field maps around the occipital pole. The locations of the V1–V3 boundaries differ slightly across the subjects, as seen by comparing the angle reversals and the V1–V3 boundary lines that were drawn based on the group-average results. This suggests that even after alignment using state-of-the-art algorithms guided by folding and areal features (MSMAll), there is residual misalignment of retinotopic maps in some subjects. A few subjects showed strikingly atypical retinotopy, such as a “forked” representation of the lower vertical meridian representation running across V2d and V3d of the group average in subjects S80 (HCP ID 198653) and S138 (HCP ID 644246) (see Figure S2 in Van Essen and Glasser, 2018 and https://balsa.wustl.edu/ZLV7).

Beyond V1–V3, several of the features we noted in the group-average results are also generally evident in the individual subjects. For example, the angle maps show a lower-field representation ventral to V3 and an upper-field representation dorsal to V3 (locations 1 and 2 in Figure 7). There are also distinct foveal representations in parietal and temporal cortex (locations 3 and 4), and pRF size gradients in ventral cortex (locations 5 and 6). Because variance explained is generally lower for individual subjects compared to the group average, there are some regions in which the group average may provide useful information that is absent in individual subjects (e.g. location 8). By visual inspection, the overall map quality appears comparable across the three subjects. Since these subjects span the central 50% of variance explained (as detailed previously), this suggests that most of the subjects in the HCP 7T Retinotopy Dataset have good data quality. Additional aspects of individual variability can be readily inspected by scrolling through polar angle and eccentricity maps for all 181 individual subjects in the downloadable Connectome Workbench ‘scene’ files (see Appendix).

#### Within-subject reliability

To quantify reliability of pRF parameters for individual subjects, we compared parameter estimates across split-halves of the data. We binned cortical grayordinates into 4 large ROIs which comprise distinct subsets of the regions in the Wang et al. atlas (Wang et al., 2015): posterior (V1–V3), dorsal (V3A/B, IPS0–5), lateral (LO-1/2, TO-1/2), and ventral (VO-1/2, PHC-1/2). We then aggregated grayordinates within each of these ROIs across subjects, and computed 2D histograms comparing parameter estimates across the two model fits (first half of each run; second half of each run).

Angle estimates were highly reliable across splits for all 4 ROIs, indicated by the high density along the diagonals (Figure 8, top row). In addition to demonstrating within-subject reliability, these histograms highlight the fact that angles near the vertical meridian (90° and 270°) are less represented than other angles, an effect observed in many prior studies (Arcaro, McMains, Singer, & Kastner, 2009; Kastner et al., 2007; Larsson & Heeger, 2006; Mackey, Winawer, & Curtis, 2017; Silver, Ress, & Heeger, 2005; Swisher et al., 2007). This effect is likely due, in part, to the fMRI signal pooling activity from neurons that represent the vertical meridian with activity from neurons that represent regions off the vertical. The eccentricity histograms (Figure 8, middle row) also show a high degree of reliability, with density highest on and near the diagonal in all 4 ROIs. Note that the dorsal ROI, while reliable, is more foveally biased than other ROIs. Nonetheless, as indicated in both the maps (Figures 4 and 7) and the reliability plots (Figure 8), the dorsal regions contain eccentricities spanning 0 to 8 degrees. Finally, the size estimates were also fairly reliable, though less so than the angle and eccentricity estimates. In agreement with the maps (Figures 4 and 7), posterior maps generally contain the smallest pRFs, with few pRF sizes larger than 3 degrees. The high reliability of pRF parameters across data splits supports the interpretation that grayordinates in not only V1–V3 but also dorsal, lateral, and ventral higher extrastriate regions exhibit spatial selectivity and not mere visual responsivity. If grayordinates responded indiscriminately to a visual stimulus presented anywhere in the visual field, pRF parameters would not exhibit such reliable tuning for specific pRF parameter values.

**Figure 8.**
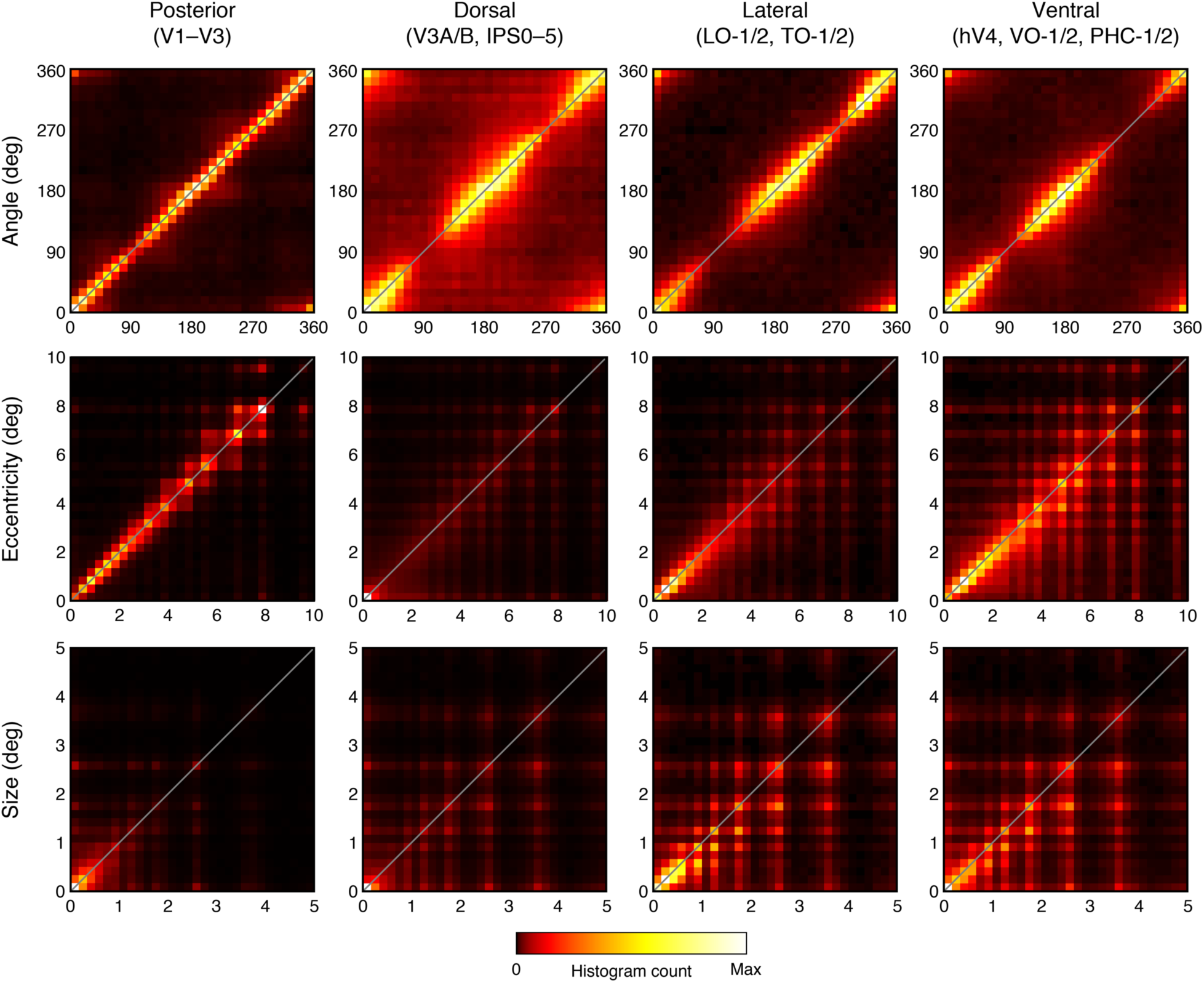
Within-subject reliability of pRF estimates. Estimates of pRF parameters were obtained for two independent splits of the data (first half of each run; second half of each run). Here, we aggregate results across all 181 individual subjects and plot 2D histograms comparing pRF parameter estimates across the two splits of the data (*x*-axis: first split; *y*-axis: second split). The depicted colormap is used to represent histogram counts from 0 to the maximum count observed in each plot. The Wang et al. atlas (Wang et al., 2015) was used to bin grayordinates into different ROIs (posterior, dorsal, lateral, ventral).

## Discussion

In this study, we have described the HCP 7T Retinotopy Dataset and the results of fitting pRF models to all 181 individual subjects as well as the group average. To facilitate quantification of model reliability, all datasets were analyzed using split-halves in addition to the full dataset. In addition to the pRF model solutions, we also make available the stimuli and analysis code used to solve the models. This allows the research community to reproduce our analyses and/or re-analyze the time-series data using different techniques. The analyses we conducted are computationally intensive, involving three independent nonlinear optimizations for each grayordinate time series. This resulted in approximately 50 million model fits that necessitated the use of a large-scale compute cluster. By providing the parameters of the solved models, we substantially lower the barrier to entry for scientists to make use of the dataset.

### Size and quality of the HCP 7T Retinotopy Dataset

Although researchers frequently collect, and occasionally make public, retinotopy datasets, such datasets have generally included no more than 20 subjects (e.g. Benson et al., 2012). To the best of our knowledge, the HCP 7T Retinotopy Dataset is the largest publicly available dataset by an order of magnitude. In addition to containing many subjects, retinotopic maps are derived from six fMRI runs (a total of 30 minutes of data), making this dataset large both in terms of number of subjects as well as amount of data per subject. Finally, the data were acquired at ultra-high magnetic field strength (7T), providing enhanced signal-to-noise ratio and high spatial and temporal resolution (1.6-mm isotropic voxels, 1-second temporal sampling using multiband data acquisition). The advantages of the dataset are clear. In individual subjects, there are reliable results, even beyond striate and extrastriate cortex. At the group level, the massive averaging of subjects reveals signals in regions of cortex (such as the inferior frontal sulcus) where conventional datasets typically have low signal that can be difficult to distinguish from noise.

### Limitations of the dataset

Though the dataset has clear value, it is also important to understand its limitations and take these into account when interpreting the data. There are several technical issues; we mention a few here, but refer to the Methods for a fuller description. The stimulus size extended to an eccentricity of 8 degrees of visual angle, and so representations of the far periphery are not well measured. Because of cortical magnification of the central visual field, robust signals are found in about half of the surface area of V1 (see Figure 5). Edge effects arise for grayordinates whose pRF centers are near the limit of the stimulus extent: these grayordinates are likely to have underestimates of pRF size and a displaced pRF center. Model solutions are somewhat discretized, reflecting the influence of the first-stage grid fit. Model solutions were constrained to have a non-negative gain factor; this may not be appropriate for studying brain regions that exhibit BOLD signal decreases in response to visual stimulation. Finally, there is an inherent limitation related to the fact that we analyzed the data using one specific pRF model with a particular code implementation. The motivation of this paper is to use established tools and models to generate a high-quality retinotopy resource, but some scientific questions will require additional modeling work. Exploring other pRF models (such as a difference-of-Gaussians pRF model (Zuiderbaan, Harvey, & Dumoulin, 2012), an anisotropic Gaussian pRF model (Merkel, Hopf, & Schoenfeld, 2018; Silson, Reynolds, Kravitz, & Baker, 2018), or pRF models with flexible shapes (Greene, Dumoulin, Harvey, & Ress, 2014; Lee, Papanikolaou, Logothetis, Smirnakis, & Keliris, 2013)) and carefully evaluating model accuracy (Kay et al., 2013a) may be important to answer such questions.

In addition to modeling choices, it is important to also consider stimulus selectivity and task effects. In particular, there may be parts of the brain that show retinotopic organization given an appropriate stimulus and task but not in all retinotopic experimental paradigms. In the experiment used for the HCP dataset, the images within the moving apertures changed 15 times per second. Cortical areas with low-pass temporal tuning might not be effectively driven by these stimuli (Liu & Wandell, 2005). Other areas, for example in the dorsal attention network, might respond strongly only when stimuli are attended (Mackey, Winawer, & Curtis, 2017); in the HCP experiment, subjects attended to the fixation location. Hence, a lack of spatial selectivity or retinotopic organization in this dataset, even with the large number of subjects, should not be taken as definitive evidence that a brain area is not spatially selective or retinotopically organized.

The neuroscientific interpretation of the pRF results must also be done carefully. Whereas in visual cortex, there is clear interpretation of pRF models in terms of visually responsive population receptive fields, in other parts of the brain, it may be possible to obtain good pRF fits but for different reasons. For example, it is possible that a cortical region indexing cognitive difficulty exhibits response increases when the stimulus is near the fovea because at these points in the experiment, the stimulus is more likely to interfere with the fixation task performed by the subjects. In such a case, the existence of a pRF model solution does not imply visually driven activity in the conventional sense.

### Group-average interpretation

The group-average datasets (S182–S184) provide a useful summary of the overall organization of visually responsive cortex. Because of the large number of subjects that are averaged together and the improved intersubject alignment methods, these datasets contain very high signal-to-noise level. However, there are several caveats to interpreting these data. Most significantly, unlike in individual subjects, the quality of pRF model fits is influenced by the quality of the alignment used to generate these averages. In particular, the quality of the fits in V1, V2, and V3 of the group average appears to be very high; this is due both to the fact that visually evoked signals in V1–V3 are particularly robust as well as the fact that these cortical areas are less variable in their locations and internal topographic structure than higher visual areas. As one moves from striate to extrastriate cortex and beyond, coverage of the visual field becomes less and less complete. The effect of averaging can be appreciated by comparing the group-average data to the individual data (see Figure 7). In the group-average data, the V1 maps sweep out nearly 180° of polar angle from the lower vertical meridian (deep red) to the upper vertical meridian (deep blue); the V3 maps do not quite reach the vertical meridians, and V3A (location 2 in the angle map) reaches only about 45° (cyan as opposed to blue). In contrast, the individual-subject maps, though noisier, often exhibit more representation of the vertical meridian in V3 and V3A (deep blue and red in the angle maps of Figure 7). Indeed, previous work has found that for most visual areas beyond V1–V3, the overlap among subjects decreases substantially (Wang et al., 2015). One analysis that could help shed light on these issues is to examine intersubject variability in pRF parameters as a function of cortical location.

### What can the HCP 7T Retinotopy Dataset be used for?

This rich dataset has a wide range of uses. It provides the basis for further analysis of other HCP data; for example, the pRF solutions for an individual subject can be used to determine visual ROIs that could then be used to analyze or validate other HCP measures. Some example applications include the following: (1) The retinotopy dataset can be used for comparison with the HCP’s multimodal parcellation (Glasser, Coalson, et al., 2016a) (see Figure 5). We have shown that the group-average results approximately agree with portions of the parcellation, but we did not compare individual-subject results to the group average parcellation or to the individual-subject parcellations that were generated using an areal classifier algorithm (Glasser et al., 2016a). (2) Identifying visual ROIs and pRF properties within the ROIs can be used in conjunction with resting-state data (Van Essen et al., 2012) to test hypotheses about how maps relate to functional connectivity. (3) The pRF model solutions can be used in conjunction with the working memory dataset (Barch et al., 2013) to study the role of visual cortex in working memory. Many more such applications (e.g. combining retinotopy with the 7T movie data) are possible.

The visuotopic mapping in the Glasser et al. (Glasser et al., 2016a) parcellation was based on resting-state fMRI correlations measured across the entire visual field representation. This enabled mapping the full extent of visuotopic areas, but does not provide explicit measurement of specific eccentricities or polar angles within each map. Hence, the current 7T retinotopic maps and the visuotopic organization derivable from resting-state data represent complementary and potentially synergistic information.

The HCP 7T Retinotopy Dataset also has a great deal of standalone value, owing to the very large number of subjects. Any examination of the relationship between anatomy and function benefits from having many subjects to characterize the extent of intersubject structure-function variability in an anatomically-normalized format. Averaging retinotopic time-series data across a large number of subjects has revealed that large swaths of cortex not typically studied by vision scientists show evidence of retinotopic organization (see Figure 4); many of these regions would not have clear signals in smaller sample sizes.

### Resolving controversies regarding retinotopic maps

Despite 25 years of measuring retinotopic maps with fMRI, a number of disagreements concerning map organization remain unresolved. For example, there are two different proposals for the organization of the V4 map and neighboring regions. The hV4/VO proposal consists of a single hV4 hemifield map on the ventral surface, with several additional hemifield maps located more anterior (Arcaro et al., 2009; McKeefry & Zeki, 1997; Wade et al., 2002; Winawer & Witthoft, 2015). The V4/V8 proposal involves a different arrangement, with the V4 map split into dorsal and ventral arms (Hansen, Kay, & Gallant, 2007; Sereno et al., 1995) and with a V8 hemifield map adjacent to ventral V4 (Hadjikhani & Tootell, 2000; Hadjikhani, Liu, Dale, Cavanagh, & Tootell, 1998; Tootell & Hadjikhani, 2001). These two different proposals are implicit in the parcellations shown in Figure 5 from Wang et al. (Wang et al., 2015) and Glasser et al. (Glasser, Coalson, et al., 2016a). There are a number of additional unresolved questions regarding map organization, including the number and arrangement of maps in the vicinity of MT (Amano, Wandell, & Dumoulin, 2009; Kolster, Peeters, & Orban, 2010). In fact, beyond the V1–V3 maps, there is likely no retinotopic map that is universally agreed upon by researchers in the field. Part of this is due to the challenge of interpreting complex spatial data; other disagreements might stem from differences across datasets, e.g., due to differences in MRI acquisition methods, stimuli, analysis approaches, and subjects. Indeed, even within V1–V3, there are unresolved questions about retinotopic organization, such as how precisely the maps align with anatomical landmarks and whether some individuals have maps that qualitatively differ from the typical pattern (see Supplemental Information in Van Essen & Glasser, 2018; https://balsa.wustl.edu/ZLV7). We believe that the HCP 7T Retinotopy Dataset provides a unique opportunity to adjudicate among competing hypotheses about the organization of retinotopic representations in the human brain. Future work, perhaps exploiting automated, objective atlas-based fitting procedures (e.g. Benson & Winawer, 2018), could help evaluate how well different proposals are supported by the data.

## Conclusion

The visual system is one of the primary functional systems of the human brain, and the resources provided in this paper represent an important step towards more fully characterizing its fundamental organization. The authors believe that the present measurements fill a critical role, both for answering novel scientific questions and for establishing baselines and hypotheses for new experiments. To this end, we have put effort into making all data and analyses fully public and well-documented, and we hope that other researchers will find this dataset enlightening and useful.

## Acknowledgements

This work was supported by the Human Connectome Project (1U54MH091657) from the 16 Institutes and Centers of the National Institutes of Health that support the NIH Blueprint for Neuroscience Research; Biotechnology Research Center (BTRC) grant P41 EB015894 from NIBIB; NINDS Institutional Center Core grant P30 NS076408; and R01 EY027401 (J.W.).

## Author Contributions

D.V.E. and K.U. planned the experiment. A.V., E.Y., and K.U. developed and optimized acquisition sequences and protocols. A.V. and K.K. designed the experiment. N.B., K.J., M.A., M.G., T.C., and K.K. analyzed the data. T.C., M.G., and D.V.E. prepared results for the BALSA database. N.B., J.W., and K.K. wrote the paper. All authors discussed the results and edited the manuscript.

## Methods

### Subjects

Complete retinotopy datasets (six fMRI runs) were acquired for a total of 181 subjects (109 females, 72 males), age 22–35, as part of the Young Adult Human Connectome Project (HCP) (https://www.humanconnectome.org/study/hcp-young-adult/data-releases). All subjects also participated in ∼4 hours of multimodal MRI data acquisition on a customized Siemens 3T ‘Connectom’ scanner at Washington University (Van Essen et al., 2013) as well as extensive behavioral and demographic assessments (Barch et al., 2013). All subjects had normal or corrected-to-normal visual acuity. The subjects include 53 pairs of genetically confirmed identical twins (106 individuals), 34 pairs of fraternal twins (68 individuals), 2 pairs of non-twin siblings (4 individuals), and 3 individuals whose twins/siblings were not included. Each subject has an assigned 6-digit HCP ID. For family structure details, researchers must apply for access to “Restricted Data” on ConnectomeDB (https://www.humanconnectome.org/study/hcp-young-adult/document/wu-minn-hcp-consortium-restricted-data-use-terms).

### Structural image acquisition and pre-preprocessing

T1-weighted (T1w) and T2-weighted (T2w) structural scans at 0.7-mm isotropic resolution were acquired at 3T and used as the anatomical substrate for the retinotopy data. White and pial cortical surfaces were reconstructed from the structural scans using the HCP Pipelines (Glasser et al., 2013). Surfaces were aligned across subjects to the HCP 32k fs_LR standard surface space using first a gentle folding-based registration ‘MSMSulc’ and then a more aggressive areal-feature-based registration ‘MSMAll’ that was driven by myelin maps, resting-state network maps, and 3T resting-state visuotopic maps (Glasser, Coalson, et al., 2016a; Robinson et al., 2018; 2014). Myelin maps were the ratio of T1w/T2w images (Glasser & Van Essen, 2011) normalized using a surface-based atlas to estimate B1+ transmit effects (Glasser et al., 2013). Note that because the MSMAll registration is based partly on measurements of resting-state networks and resting-state-based estimates of visuotopic organization (Glasser et al., 2016a), alignment of pRF solutions across individuals is likely to be improved relative to alignment based on cortical folding alone. Subcortical volume data were aligned to MNI space using FNIRT nonlinear volume-based registration based on T1w image intensities (Glasser et al., 2013).

### fMRI acquisition and pre-processing

Full details on data acquisition and pre-processing are provided elsewhere (Glasser et al., 2013; Vu et al., 2016). In brief, fMRI data were collected at the Center for Magnetic Resonance Research at the University of Minnesota using a Siemens 7T Magnetom actively shielded scanner and a 32-channel receiver coil array with a single channel transmit coil (Nova Medical, Wilmington, MA). Whole-brain fMRI data were collected at a resolution of 1.6-mm isotropic and 1-s TR (multiband acceleration 5, in-plane acceleration 2, 85 slices). The data were processed using the HCP pipelines (Glasser et al., 2013) that correct for head motion and EPI spatial distortion and bring the fMRI data into alignment with the HCP standard surface space as described above. (Note that the data used here reflect the correct phase-encode directions in the EPI undistortion procedure, unlike an early pre-2018 release of the data.) The data produced by the pipeline are in the CIFTI format, which consists of 91,282 grayordinates that cover both cortical and subcortical brain regions with approximately 2-mm spatial resolution. (Higher-resolution CIFTI outputs are also available, consisting of 170,494 grayordinates with approximately 1.6-mm spatial resolution. Only the 2-mm CIFTI data are used in this paper.) The fMRI data were also denoised for spatially specific structured noise using multi-run sICA+FIX (Glasser et al., 2018; Griffanti et al., 2014; Salimi-Khorshidi et al., 2014). Differences in slice timing were not corrected since the fast multiband acquisition makes such corrections less important (though slices may differ by as much as 1 s). The dimensions of the pre-processed data are 181 subjects × 91,282 grayordinates × 6 runs × 300 time points. These pre-processed data are available from ConnectomeDB (https://db.humanconnectome.org/).

The HCP’s methods of pre-processing are designed to maximize alignment of cortical areas across subjects while minimizing blurring of the data in individuals or groups (Coalson, Van Essen, & Glasser, 2018; Glasser, Smith, et al., 2016b). HCP-style pre-processing includes correction of distortions in all MRI images so that the images represent the physical space of the subject, registration across modalities and correction for motion within fMRI and DWI scans, and bringing cortical data onto surface meshes and subcortical data in gray-matter parcels (Glasser et al., 2013). Data are aligned on the surface using MSMAll areal-feature-based registration, which uses myelin maps and resting-state data to more accurately align cortical areas than a standard folding-based registration (Glasser, Coalson, et al., 2016a; Robinson et al., 2014; 2018). Data from subcortical areas are aligned using FNIRT nonlinear volume-based registration. Care was taken to minimize the number of interpolations and to use interpolation methods like splines that minimize the blurring effects of interpolation.

Despite efforts in careful pre-processing methods, imperfections inevitably remain in the data. Partial volume effects and other types of blurring are inherent in data acquisition and cannot be easily removed through pre-processing methods. Thus, care must be taken in interpretation where different tissue types are in close proximity. For example, it appears that the dorsal rim of the cerebellum may exhibit visually responsive signals that likely originate, in part, from nearby locations in cortex.

### Stimuli

Retinotopic mapping stimuli were constructed by creating slowly moving apertures and placing a dynamic colorful texture within the apertures. Apertures and textures were generated at a resolution of 768 pixels × 768 pixels, and were constrained to a circular region with diameter 16.0°. The display was uniform gray beyond the circular region.

#### Texture design

To elicit strong neural responses in high-level visual areas (while also driving responses in early visual areas), we designed a texture composed of colorful visual objects. The texture was constructed by taking objects from Kriegeskorte et al., 2008, preparing these objects at multiple scales, and placing the objects on an achromatic pink-noise background. One hundred (100) distinct texture images were generated. To generate a texture image, we first created an achromatic pink-noise (1/*f* amplitude spectrum) background. Then, starting at the largest scale and proceeding to smaller scales, objects were randomly selected and placed at random positions in the image, potentially occluding objects already placed, similar to a ‘dead leaves’ tessellation. There were seven different scales. The object sizes associated with the seven scales were 350 (7.3°), 247 (5.1°), 175 (3.6°), 124 (2.6°), 88 (1.8°), 62 (1.3°), and 44 (0.9°) pixels (decreasing by a factor of v2), and the numbers of objects at each scale were 1, 2, 4, 8, 16, 32, and 64 (increasing by a factor of 2). Textures were updated at a rate of 15 Hz (details below). Pilot experiments confirmed that compared to conventional checkerboard patterns, the object texture produces larger BOLD responses and improves test-retest reliability of retinotopic estimates.

#### Aperture design

The experiment consisted of six runs in which three different types of apertures were presented (wedges, rings, bars). Apertures moved slowly across the visual field, and were occasionally interrupted by blank periods in order to help distinguish between non-visual responses and responses from neurons with very large receptive fields (Dumoulin & Wandell, 2008). Each run lasted 300.0 s. The order of runs was RETCCW, RETCW, RETEXP, RETCON, RETBAR1, and RETBAR2, and are described below:

- RETCCW consisted of a 22-s blank period, 8 cycles of a 90° wedge rotating counter-clockwise with a period of 32 s, and a 22-s blank period. The duty cycle for a given point in the visual field was 25% (8 s of 32 s).
- RETCW was the same as RETCCW except that the wedge rotated clockwise.
- RETEXP consisted of a 22-s blank period, 8 cycles of a ring expanding away from the center of the screen with a period of 32 s, and a 22-s blank period. The last 4 s of each 32-s period was blank (thus helping distinguish foveal and peripheral responses). Ring size increased linearly with eccentricity. The duty cycle for a given point in the visual field was 19% (6 s of 32 s).
- RETCON was the same as RETEXP except that the ring contracted towards the center of the screen.
- RETBAR1 and RETBAR2 were identical, and consisted of a 16-s blank period, 4 bar movements lasting 32 s each (RIGHT, UP, LEFT, DOWN), a 12-s blank period, 4 bar movements lasting 32 s each (UPPER-RIGHT, UPPER-LEFT, LOWER-LEFT, LOWER-RIGHT), and a 16-s blank period. The capitalized term indicates the direction of bar movement. The last 4 s of each 32-s bar movement was blank (thus, the bar traversed the visual field in 28 s). The width of the bar was 1/8 of the full stimulus extent. The duty cycle for a given point in the visual field was 10% (3.11 s of 32 s).

Apertures were animated at a rate of 15 Hz, and each aperture was anti-aliased to minimize discretization effects. On each aperture update, one of the 100 texture images was randomly selected (under the constraint that the same texture image is not presented consecutively) and presented within the confines of the aperture (using the continuous values of the aperture as opacity values). Each run consisted of 300 s × 15 Hz = 4500 stimulus frames.

### Experimental design and task

A small semi-transparent dot (0.3° × 0.3°) at the center of the display was present throughout the experiment. The color of the central dot switched randomly to one of three colors (black, white, or red) every 1–5 s. Subjects were instructed to maintain fixation on the dot and to press a button whenever the color of the dot changed. The purpose of the task was to encourage fixation and allocation of attention to the center of the display. To further aid fixation, a semi-transparent fixation grid was superimposed on the display throughout the experiment (Schira, Tyler, Breakspear, & Spehar, 2009).

Stimuli were presented using an NEC NP4000 projector. The projected image was focused onto a backprojection screen, and subjects viewed this screen via a mirror mounted on the RF coil. The projector operated at a resolution of 1024 × 768 @ 60 Hz. A Macintosh computer controlled stimulus presentation using code based on the Psychophysics Toolbox (Brainard, 1997; Pelli, 1997). Behavioral responses were recorded using a Curdes FORP button box. Eye-tracking was performed using an EyeLink 1000 system (SR Research, Mississauga, Ontario, Canada). Eye-tracking data are available on ConnectomeDB for most subjects, but we caution that the quality of the data is variable due to obstructions within the head coil. Eye-tracking data were not used in this paper.

The viewing distance to the backprojection screen was 101.5 cm, and the full stimulus extent (i.e. diameter of the circle within which apertures are shown) was 28.5 cm, yielding a total stimulus size of 16.0°. However, due to variations in subject setup, these numbers should be considered approximate. Furthermore, due to the confines of the MRI environment, some subjects were unable to see the very top and very bottom of the stimuli (approximately 1° at each end). This should be taken into account when interpreting the fMRI results.

### pRF analysis

#### Population receptive field (pRF) model

We analyzed the time-series data of each grayordinate using a pRF model called the Compressive Spatial Summation model (Kay et al., 2013a). This model is implemented in a MATLAB toolbox called analyzePRF (http://cvnlab.net/analyzePRF/); to analyze the HCP 7T Retinotopy Dataset, we modified the implementation and archived the resulting code on the Open Science Framework web site (https://osf.io/bw9ec/). The model predicts the fMRI time series as the sum of a stimulus-related time series and a baseline time series. The stimulus-related time series is obtained by computing the dot product between the stimulus apertures and a 2D isotropic Gaussian, applying a static power-law nonlinearity, scaling the result by a gain factor, and then convolving the result with a canonical hemodynamic response function (HRF). This can be expressed formally as *r*(*t*) = (*g* × (*S*(*t*)•*G*)^*n*^) * *h*(*t*) where *r*(*t*) is the predicted stimulus-related time series, *g* is a gain parameter, *S*(*t*) is the stimulus aperture at time *t, G* is the 2D isotropic Gaussian, *n* is an exponent parameter, and *h*(*t*) is a canonical HRF. This time series characterizes BOLD modulations driven by the stimulus. The baseline time series is obtained by computing a weighted sum of low-order polynomial terms (constant, linear, quadratic, etc.). This time series characterizes the baseline BOLD signal level, i.e., the MR signal intensity that is present in the absence of the stimulus. The canonical HRF was obtained by taking an ideal impulse response determined in a previous study (Kay et al., 2013b) (spm_hrf(0.1,[6.68 14.66 1.82 3.15 3.08 0.1 48.9]) from SPM), convolving the impulse response to predict the response to a 1-s stimulus, resampling to a 1-s sampling rate using cubic interpolation, and normalizing the result to have a peak amplitude of 1. Note that this HRF was used for all grayordinates in all subjects.

The model yields several parameters of interest: two parameters (*x, y*) that indicate the position of the Gaussian, a parameter (*σ*) that indicates the standard deviation of the Gaussian, a parameter (*n*) that indicates the exponent of the power-law nonlinearity, and a parameter (*g*) that indicates the overall gain of the predicted responses in raw scanner units. In pilot analyses, we found that the experimental paradigm used here generally does not provide enough statistical power to estimate the exponent parameter reliably. Thus, we did not attempt to estimate this parameter but instead fixed the value of *n* to 0.05, which is representative of the typical values that we observed in the dataset. Note that a compressive exponent (like 0.05) has the effect of making responses tolerant to changes in the position and size of the stimulus (Kay et al., 2013a); intuitively, with a compressive exponent, stimuli presented anywhere within the pRF tend to drive responses equally strongly.

*pRF size*

The spatial selectivity of a grayordinate depends on both the size of the Gaussian and the sub-additive exponent. This relationship can be made explicit by considering the predicted response to point stimuli at different locations in the visual field. As described previously (Kay et al., 2013a), these responses can be expressed as:

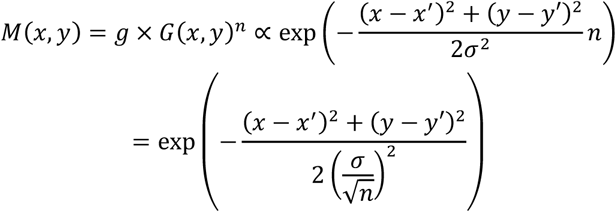

where *M* indicates responses to the different point stimuli, *g* is the pRF gain, *G* is the 2D isotropic Gaussian parameterized by position (*x’, y’*) and standard deviation (σ), and *n* is the sub-additive exponent. Note that these responses, *M*, follow the form of a new Gaussian. We define pRF size as the standard deviation of this new Gaussian:

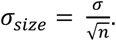

Because we fix *n* at 0.05, the pRF size is ∼4.5 times larger than the standard deviation of the Gaussian *G*. Throughout the text, all references to pRF size reflect σ_size._ Hence, the reported pRF size already accounts for the effect of the power-law nonlinearity. In summary, pRF size can be interpreted as one standard deviation of a 2D Gaussian that characterizes how a grayordinate responds to point stimuli in the visual field.

#### Stimulus pre-processing

Prior to model fitting, we performed pre-processing of the stimulus apertures. The original resolution of the apertures in each run is 768 pixels × 768 pixels × 4500 frames. Aperture values range between 0 and 1, where 0 indicates the absence of the texture image and 1 indicates the presence of the texture image. To reduce computational burden, we resized the apertures to 200 pixels × 200 pixels. Then, to match the temporal resolution of the stimulus to the temporal resolution of the fMRI data, we averaged consecutive groups of 15 frames. This yielded a final stimulus resolution of 200 pixels × 200 pixels × 300 frames. Model fitting was performed in pixel units, and model parameters were *post hoc* converted from pixel units to degrees by multiplying by a scaling factor of 16.0° / 200 pixels.

#### Model fitting

In pilot analyses of the fMRI data, we noticed a high propensity for local minima in model solutions. To reduce inaccuracies and biases due to local minima, we designed the following fitting approach. We first performed a grid fit in which a range of parameter combinations were evaluated. We densely sampled parameter space using 25 nonlinearly spaced eccentricity values between 0 and 16 degrees (0, 0.04, 0.09, 0.16, 0.22, 0.33, 0.43, 0.58, 0.73, 0.95, 1.2, 1.5, 1.8, 2.2, 2.7, 3.3, 3.9, 4.7, 5.6, 6.8, 8.0, 9.6, 11.3, 13.7, and 16 degrees), 32 angle values between 0 and 360 degrees (0, 11.25, 22.5,…, and 348.75 degrees), and 13 size values on a log scale between 1 and 128 pixels (equivalent to 0.08, 0.11, 0.16, 0.23, 0.32, 0.45, 0.64, 0.91, 1.3, 1.8, 2.6, 3.6, 5.1, 7.2, and 10.2 degrees), yielding 25 × 32 × 13 = 10,400 parameter combinations. The combination yielding the optimal fit (in a least-squares sense) to the data was identified. We then used this parameter combination as the initial seed in a nonlinear optimization procedure (MATLAB Optimization Toolbox, Levenberg-Marquardt algorithm). For this initial seed, the gain parameter was changed to 75% of its discovered (optimal) value to allow room for adjustment in the optimization. Also, the gain parameter was restricted to be non-negative to constrain the space of fits to solutions that predict positive BOLD responses to stimulation. Note that no spatial constraints are incorporated into the model fitting process (e.g., smoothing, priors on expected parameter values, etc.). Thus, parameter estimates for grayordinates are independent, thereby maximizing resolution and minimizing bias.

We fit the pRF model not only to the data from each subject (S1–S181), but also to the data from three group-average pseudo-subjects, which were constructed by averaging time-series data across subjects. One group average is the result of averaging all 181 subjects (S184); the second group average is the result of averaging a randomly chosen half of the subjects (S182); and the third group average is the result of averaging the other half of the subjects (S183). For each individual subject and each group average, we performed three separate model fits: one fit uses all six runs, a second fit uses only the first half of each of the six runs, and the third fit uses only the second half of each of the six runs. The rationale for these fits is that the first fit provides the best estimate of model parameters, whereas the second and third fits can be used to assess the reliability of parameter estimates.

Each fit produces six quantities of interest: pRF angle, pRF eccentricity, pRF size (calculated as *σ*/v*n*), pRF gain, percentage of variance explained, and mean signal intensity (calculated as the mean of all time points). The dimensions of the final results are (181+3) subjects × 91,282 grayordinates × 3 model fits × 6 quantities (see Figure 2).

#### Surface visualization

The pre-processed time-series data (CIFTI format) reflect MSMAll-alignment of individual subjects to the fs_LR surface (Glasser et al., 2013). The pRF model solutions are obtained by fitting each CIFTI grayordinate independently; thus, there are no additional spatial transformations applied. In this paper, we visualize pRF model solutions for cortical grayordinates by mapping from CIFTI space to *fsaverage* space using nearest-neighbor interpolation and then using orthographic projection to visualize the *fsaverage* surface (see Figure 3). The underlay for the group-average results is the thresholded *fsaverage* curvature, whereas the underlay for individual-subject results is the curvature obtained from individual subjects. An alternative to the *fsaverage* curvature is to compute the average curvature of the 181 subjects, which is useful for maintaining the true relationship between the retinotopic features and folding features in the data (e.g. as in Glasser et al. 2016a); this average curvature is the underlay provided in the downloadable BALSA datasets (https://balsa.wustl.edu/study/show/9Zkk).

Group-average maps in Figures 3–6 are thresholded at 9.8% variance explained. This threshold was determined by fitting a Gaussian Mixture Model with two Gaussians to the distribution of variance explained values across grayordinates in the group-average data and then identifying the value at which the posterior probability switches from the Gaussian with smaller mean to the Gaussian with larger mean. The interpretation of this procedure is that the Gaussian with smaller mean likely reflects noise (grayordinates that are not visually responsive), the Gaussian with larger mean likely reflects signal (grayordinates that are visually responsive), and values above the threshold are more likely to reflect signal than noise. The same procedure was performed for the distribution of variance explained values across grayordinates in individual-subject data, and this yielded a threshold of 2.2% variance explained. We used this more liberal threshold for the individual-subject maps in Figure 7.

### Timing and behavioral analysis

Stimulus timing, scanner timing, and button presses were logged in a behavioral file for each run. Behavioral files are missing for a small fraction of the runs (17 of 1086) and button presses were not detected for one of the subjects (S152, HCP ID 782561). Analysis of stimulus timing indicates that run durations were highly reliable: the central 95% of run durations lie within the range [299.974 s, 299.982 s]. Analysis of scanner timing indicates that synchronization of the stimulus computer and the scanner was robust, with the exception of two runs in which scanner acquisition may have started 1 s (1 TR) too early relative to the stimulus.

To quantify behavioral performance, we calculated, for each run, (*A*–*B*)/*C* × 100, where *A* indicates the number of successful detections of color changes (defined as the existence of a button press within 1 s of a color change), *B* indicates the number of extraneous button presses, and *C* indicates the total number of color changes. We then averaged this performance value across the six runs. Behavioral performance was quite good overall: the interquartile range (central 50%) of performance values across subjects is [91.6%, 97.8%].

## Supplementary Material

**Supplementary Movies 1–12.**
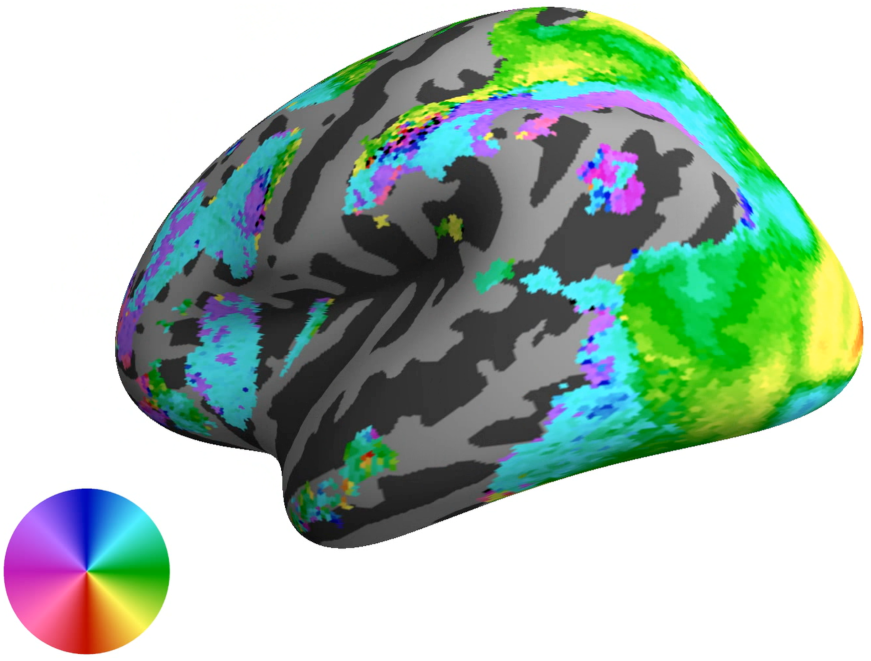
pRF results on dynamic rotating cortical surfaces. Each movie shows group-average pRF model solutions (S184) on the inflated *fsaverage* surface. The format is the same as in Figure 4. There are a total of 2 hemispheres × 6 maps (curvature, angle, eccentricity, zoomed eccentricity, pRF size, variance explained) = 12 movies. The movies are accessible at the OSF web site (https://osf.io/bw9ec/).

---

**Supplementary Figure 1.**
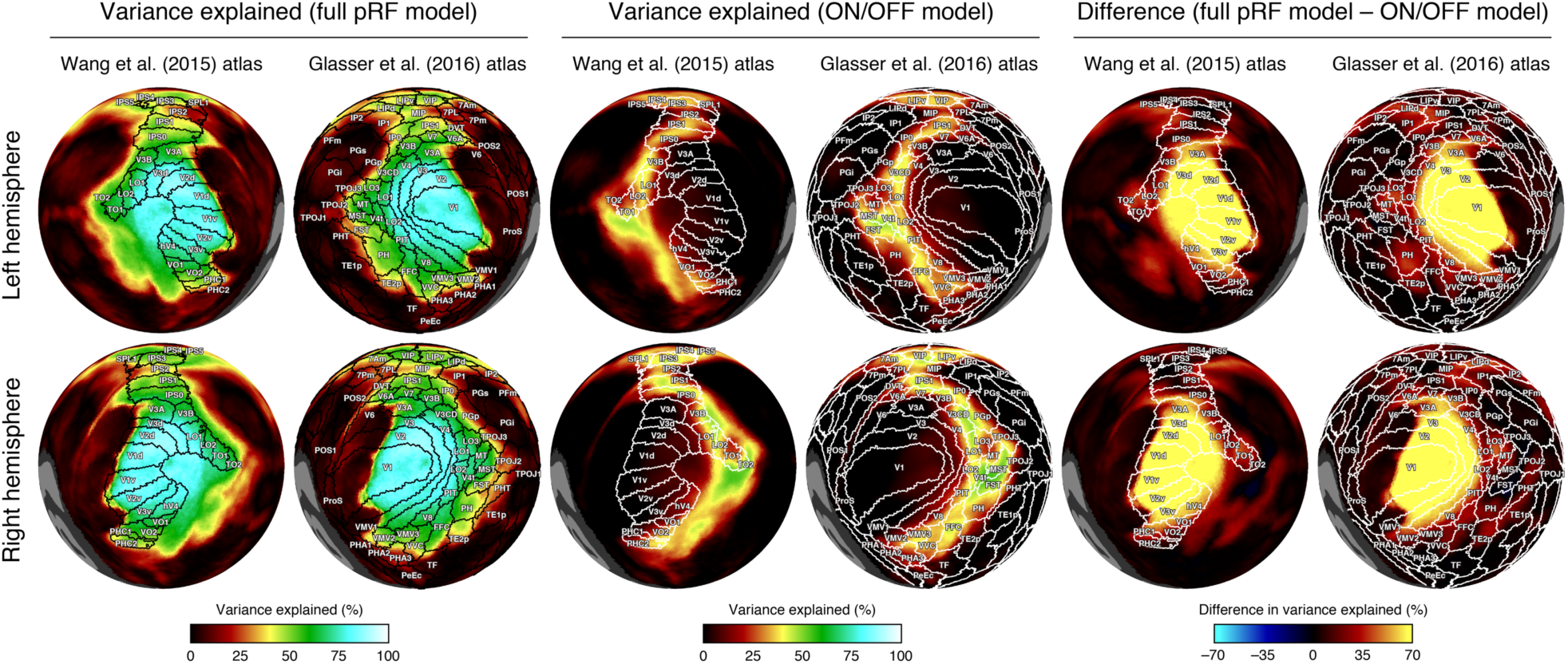
Comparison of full pRF model to a simple ON/OFF model. It is possible that the pRF model, when fitted to a grayordinate, successfully explains variance in the time-series data, even if the grayordinate possesses little or no spatial selectivity. To assess this issue, we constructed a simple model that is sensitive only to the presence or absence of the retinotopic mapping stimulus. In this model (code available on the OSF web site), the stimulus-related time-series is computed as the convolution of the canonical HRF with a time-series whose value is 1 when a stimulus is present in the visual field and 0 when a stimulus is absent. This ON/OFF model is an asymptotic case of the pRF model as pRF size approaches infinity. This figure depicts for the group average (S184), the variance explained by the full pRF model, the variance explained by the ON/OFF model, and the difference between the two. As expected, the ON/OFF model fares poorly in early visual areas (V1–V3) but performs increasingly well in later visual areas. This is consistent with the fact that later visual areas have increasingly large pRFs. However, even in lateral (e.g. LO-1/2, TO-1/2), ventral (e.g. hV4, VO-1/2), and parietal regions (e.g. MIP, IP0, IP1), the full pRF model explains substantially more variance than the ON/OFF model. This indicates that these regions are not only visually responsive but also exhibit some degree of spatial selectivity.

